# Seven-up acts in neuroblasts to specify adult central complex neuron identity and initiate neuroblast decommissioning

**DOI:** 10.1101/2023.11.02.565340

**Authors:** Noah R. Dillon, Laurina Manning, Keiko Hirono, Chris Q. Doe

## Abstract

An open question in neurobiology is how diverse neuron cell types are generated from a small number of neural stem cells. In the *Drosophila* larval central brain, there are eight bilateral Type 2 neuroblast (T2NB) lineages that express a suite of early temporal factors followed by a different set of late temporal factors and generate the majority of the central complex (CX) neurons. The early-to-late switch is triggered by the orphan nuclear hormone receptor Seven-up (Svp), yet little is known about this Svp-dependent switch in specifying CX neuron identities. Here, we (i) birthdate the CX neurons P-EN and P-FN (early and late, respectively); (ii) show that Svp is transiently expressed in all early T2NBs; and (iii) show that loss of Svp expands the population of early born P-EN neurons at the expense of late born P-FN neurons. Furthermore, in the absence of Svp, T2NBs fail decommissioning and abnormally extend their lineage into week-old adults. We conclude that Svp is required to specify CX neuron identity, as well as to initiate T2NB decommissioning.

**Summary:** Seven-up acts in Type 2 neuroblasts to specify adult central complex columnar neuron identity and to initiate neuroblast decommissioning.

## Introduction

Developing a complex brain requires neural stem cells to generate both a large and diverse set of neuron subtypes. *Drosophila* neuroblasts (NBs) are neural stem cells which generate neuronal diversity through the initiation of spatial patterning that establishes lineage identities (Skeath and Thor, 2003; Erclick et al., 2017) and subsequent temporal patterning within a lineage to produce unique neuron subtypes (Doe, 2017; El-Danaf et al., 2022). These processes have been extensively studied in embryonic ventral nerve cord NB lineages (Isshiki et al., 2001; Novotny et al., 2002; Pearson and Doe 2003; Tran and Doe 2008; Moris-Sanz et al., 2014; Grosskortenhaus et al., 2005, 2006), but the role of temporal patterning mechanisms in larval central brain NB lineages remains understudied.

The larval central brain contains ∼100 NBs per hemibrain with spatially stereotyped lineages (Pereanu and Hartenstein, 2006). Type 2 neuroblast (T2NB) lineages are generated by eight T2NBs in each hemibrain, with unique spatially defined lineage identities (Pereanu and Hartenstein, 2006; Bello et al., 2008; Boone and Doe, 2008; Bowman et al., 2008). Lineage tracing shows that each T2NB lineage (DM1-6 and DL1-2) produces neurons with lineage-specific morphology (Riebli et., al 2013; Yang et al., 2013; Andrade et al., 2019). T2NBs generate a series of intermediate neural progenitors (INPs), and each INP produces 4-6 ganglion mother cells, which each divide to produce two post-mitotic neurons/glia (Fig. 1A). Notably, the T2NB division pattern is analogous to outer subventricular zone lineages in the primate cortex (Holguera and Desplan, 2018).

**Figure 1.**
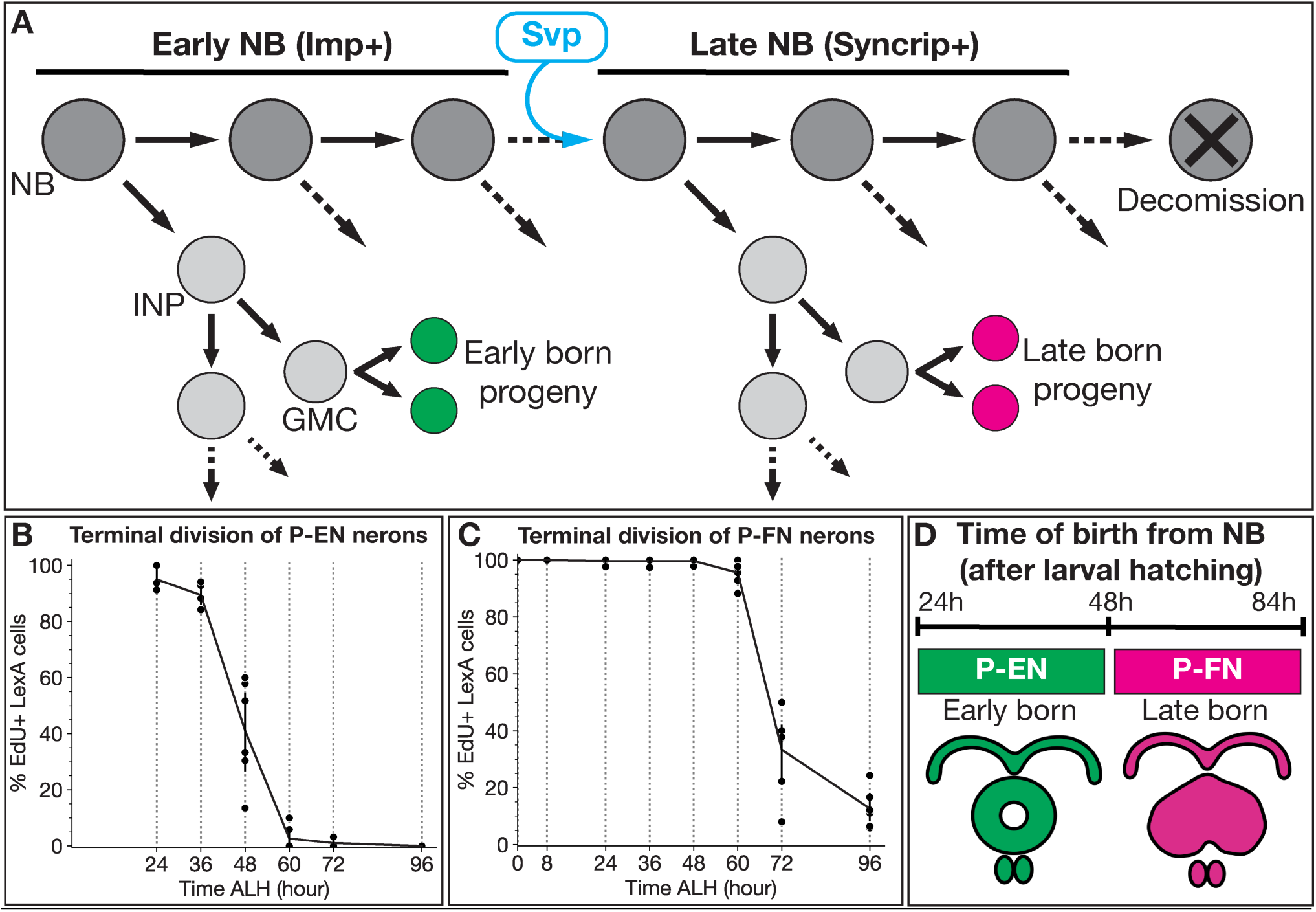
Columnar neuron subtypes are born from larval T2NBs at distinct temporal windows. (A) T2NBs express Imp during early larval stages and transition to Syncrip expression in late larvae due to Svp expression (Ren et al., 2017). All non-mushroom body NBs enter decommissioning in the early pupae (Ito and Hotta, 1992; Maurange et al., 2008; Siegrist et al., 2010; Yang et al., 2017). (B-C) Terminal division by EdU drop out for P-EN neurons (B) and P-FN neurons (C) shown by percent neurons labeled by EdU. Each dot represents one adult brain. For both P-EN and P-FN neurons at each timepoint, n = 3-7 brains. (E) Summary of P-EN and P-FN birth windows from larval T2NBs.

The T2NBs have been shown to express several genes in a temporal-specific manner during larval stages. The IGF-II mRNA-binding protein (Imp) is expressed in a temporal gradient in T2NBs, and other lineages, with high levels of Imp in early NBs and low levels in late NBs; this in inverse to the low-to-high temporal gradient of the RNA-binding protein Syncrip (Liu et al., 2015; Ren et al., 2017; Syed et al., 2017). Previous work has identified the orphan nuclear hormone receptor Seven-up (Svp) as a switching factor to initiate this Imp-to-Syncrip transition within T2NBs (Fig. 1A) (Ren et al., 2017; Syed et al., 2017). Similarly, Svp in ventral nerve cord NB lineages is required to switch Type 1 NBs from producing early born neuron fates to late born fates (Kanai et al., 2005; Mettler et al., 2006; Benito-Sipos et al., 2011; Kowhi et al., 2011). Although Svp is required to switch from early to late temporal gene expression in T2NBs, it is unknown whether this has any effect on the specification of post-mitotic neurons.

T2NB lineages generate the majority of the central complex (CX) of the *Drosophila* adult brain, with recent connectomes showing the CX containing hundreds of morphologically distinct neuron subtypes (Franconville et al., 2018; Hulse et al., 2021). One group of interest is columnar neurons, which are members of neural circuits responsible for locomotion and spatial navigation behaviors that are integrated in the CX (Giraldo et al., 2018; Green et al., 2019; Turner-Evans et al., 2020); thus, understanding the development of T2NB lineages may shed light on how complex circuits form and drive behavior.

Here, we characterize the function of Svp in specifying post-mitotic neuron identities within T2NB lineages. We find the birth windows of P-EN and P-FN columnar neurons are from early and late T2NBs respectively. We find that Svp is transiently and asynchronously expressed in all eight larval T2NB lineages. We used CRISPR/Cas9 knockout lines to remove *svp* specifically from T2NB lineages and show that Svp is required for the late born P-FN fate while restricting the early born P-EN fate. Surprisingly, we discover a novel role for Svp in terminating T2NB neurogenesis. We propose that Svp is essential for the early-to-late transition in T2NB lineages, and that these changes propagate down to the level of altered identity in post-mitotic neurons. Finally, we document a novel function of Svp in promoting NB decommissioning.

## Results

### Columnar neurons P-EN and P-FN are born from larval T2NBs in different temporal windows

Previous birth dating analysis of columnar neuron subtypes only assayed a subset of neurons, for example only 50-65% of the P-FN and P-EN neurons were birth dated (Sullivan et al., 2019). To obtain more complete coverage, we developed a 5-ethynyl-2’-deoxyuridine (EdU) drop out approach to determine the temporal birth window for P-EN and P-FN neurons. Briefly, larvae were fed EdU, a thymidine analog that incorporates into DNA during DNA synthesis. We initiated EdU feeding at different timepoints of larval life and maintained EdU feeding until pupation. By imaging adult brains from larvae fed EdU, we characterized the time of P-EN and P-FN terminal division, as neurons drop out from EdU+ to EdU- at progressively later time points of EdU introduction (Fig. 1B,C).

To account for the differences in the timing of neuron terminal division and the time its parental INP was born from the NB, we took advantage of previous cell cycle data for NBs, INPs, and ganglion mother cells (Homem et al., 2013). Both P-EN and P-FN neurons are derived from young INPs (Sullivan et al., 2019), and neurons from young INPs are born from their parental NB ∼12h prior to terminal division (Homem et al., 2013). Therefore, we subtracted 12h to reveal the time of origin from the T2NB. We conclude that P-EN neurons are born from T2NBs between 24h-48h ALH, whereas P-FN neurons are born between 48h-84h ALH (Fig. 1D). Thus, P-EN and P-FN neurons are derived from the same DM1-DM4 T2NB lineages (Yang et al., 2013) and the same young INP lineage (Sullivan et al., 2019), but differ in the timing of their birth window from the T2NB. Next, we use these early and late born neuron identities to determine whether Svp -- known for its role in regulating temporal gene expression in T2NBs (Ren et al., 2017; Syed et al., 2017) -- is also required for proper specification of post-mitotic neurons.

### Svp is expressed transiently and asynchronously in all larval T2NB lineages between 18-24h ALH

Before assaying neuron identity following Svp knockout, we assayed for Svp protein expression in each of the eight T2NB lineages, primarily to confirm expression in the DM1-DM4 T2NB lineages known to generate the columnar neurons (Riebli et., al 2013; Yang et al., 2013; Andrade et al., 2019). T2NBs were identified by the co-expression of Pnt-Gal4 driving UAS membrane-bound GFP and the pan-neuroblast marker Deadpan in cells ≥5 μm in diameter. Individual T2NB lineages were identified based on spatial position of the T2NBs within the brain lobes (Pereanu and Hartenstein, 2006; Izergina et al., 2009). We found that Svp was transiently expressed in all eight lineages with peak occurrence of expression in DM1-3 at 24h ALH (Fig. 2A-C’,I) and for lineages DM4-6 and DL1-2 at 18h (Fig. 2D-H’,I). Svp protein was restricted to the T2NB and absent in its progeny (Fig. 2A-H’). We saw a similar trend of Svp mRNA expression in all early T2NB lineages (Fig. S1). We conclude that Svp is transiently expressed in all T2NB lineages.

**Figure 2.**
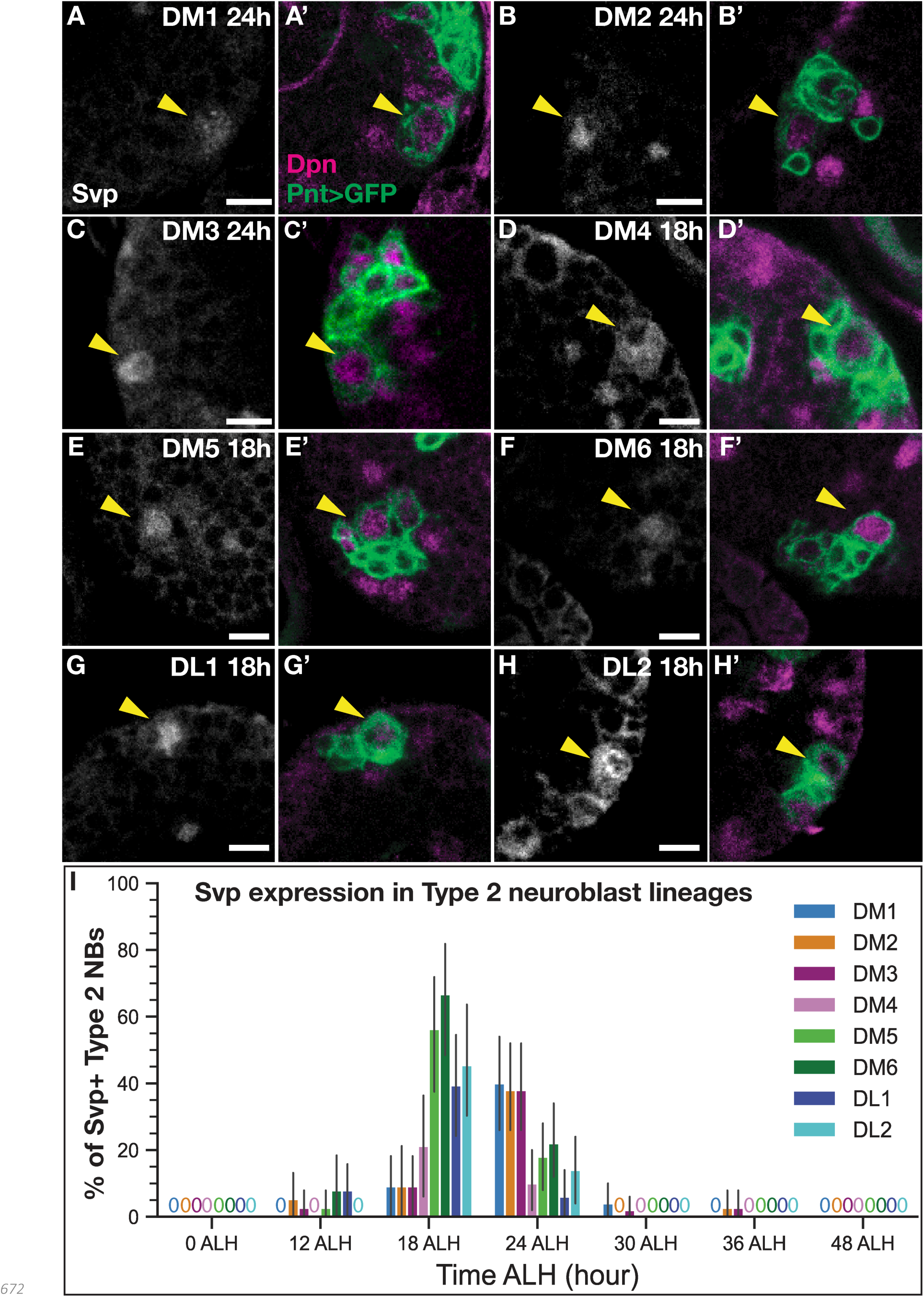
Svp is expressed early in all larval T2NB lineages. (A-C’) Svp is expressed at 24h after larval hatching (ALH) in T2NB lineages DM1-3. (D-H’) Svp is expressed at 18h ALH in T2NB lineages DM4-6 and DL1-DL2. (A-H) In all images, Svp is in white and T2NBs identified with Pnt- Gal4>GFP and Dpn. Yellow arrowhead, T2NB. (I) Quantification of Svp expression in T2NBs across 0h-48h ALH shown as a bar plot with 95% confidence interval. For each lineage, 0 ALH, n = 34; 12 ALH, n = 38; 18h ALH, n = 33; 24h ALH, n = 50; 30h ALH, n = 50; 36h ALH, n = 38; 48h ALH, n = 49 (DM lineages) or 42 (DL lineages) lobes. Scale bars: 5 μm.

We performed Svp loss-of-function with the goal of making the most complete loss- and gain-of-function alterations to the T2NB temporal factor cascade as possible, thereby increasing the chance of seeing changes in P-EN and P-FN neuron identities. We chose to knockout *svp* specifically in T2NB lineages, which has been shown to extend the expression of early NB factors (e.g. Imp, Chinmo) at the expense of late NB factors (e.g. Syncrip, EcRB1, Broad, E93) (Ren et al., 2017; Syed et al., 2017); although changes in neuronal morphology were detected, adult post-mitotic neuron molecular identity was not assayed in these experiments. We hypothesize that if early NB factors play a role in neuronal specification, then loss of Svp should result in ectopic P-EN neurons due to failure to switch to late temporal factors, and conversely, Svp knockout should reduce or eliminate late born identities such as P-FN neurons.

We generated Svp knockouts specifically in T2NB lineages. We used two independent CRISPR/Cas9 lines (Port et al., 2020) each with two Svp-specific sgRNAs to knockout *svp* in T2NBs. To determine the efficiency of our knockouts, we tested these lines for the loss of Svp expression and recapitulation of the loss of the late T2NB expression of E93 (Syed et al., 2017). We find that our knockout lines significantly reduce the occurrence of Svp expression in T2NBs but with a minority of escaper NBs that have expression levels indistinguishable from wild-type (Fig. S2A-D); these are likely 1-2 T2NB lineages where CRISPR/Cas9-mediated *svp* knockout did not occur. Thus, our Svp knockouts had “all or none” effects on Svp expression. T2NBs that exhibit loss of Svp also show a loss of E93 expression, as previously observed (Syed et al., 2017), further validating the Svp knockouts (Fig. S2E-H). We conclude that our CRISPR/Cas9 lines effectively knockout *svp* completely within the majority of T2NB lineages with only a few escapers that show wild-type levels of Svp expression. We conclude that these knockout lines can be used to test the role of Svp in specifying adult columnar neuron identity.

### Cut expression distinguishes molecular identities of adult P-EN neurons from P-FN neurons

P-EN and P-FN neurons were first characterized based on their distinct axon projections into CX neuropils (P-EN sends axons to the EB; P-FN sends axons to the FB) (Wolff et al., 2015). Yet to date no molecular markers have been reported to distinguish these neurons, except for the LexA lines we use here. To address this, we used a single-cell RNA-sequencing atlas of adult T2NB-derived neurons to identify novel molecular markers for P-EN and P-FN neurons (D. Epiney, G. Morales, NRD, S-L. Lai, and CQD; in preparation). We identified that the homeodomain transcription factor Cut was expressed in adult P-EN neurons but not in P-FN neurons (Fig. 3A-B’’,C), whereas Svp was in neither adult neuron type (Fig. S3A-B’’,C) and Runt was in both neuron types, as previously reported (Sullivan et al., 2017) (Fig. 3A’,B’; Fig. S2A-B’). We conclude that Cut distinguishes the molecular identities of adult P-EN and P-FN neurons, and we use this marker in combination with subtype-specific LexA-driven V5 and Runt expression in our analysis of the Svp knockout phenotypes.

**Figure 3.**
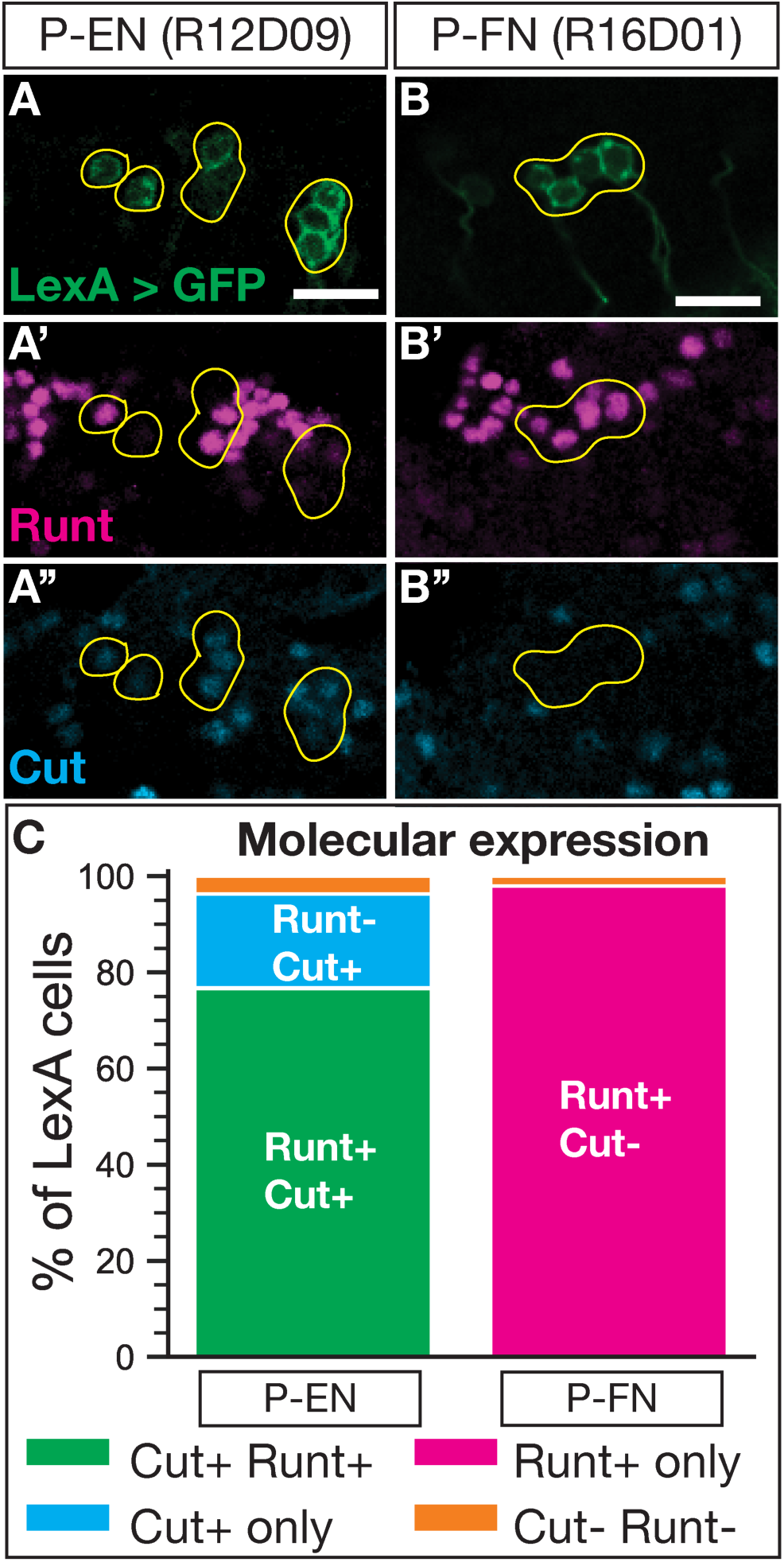
Cut expression distinguishes P-EN and P-FN molecular identities. (A-A’’) P-EN neurons labeled by LexA driver co-express Runt and Cut. (B-B’’) P-FN neurons labeled by LexA driver express Runt but not Cut. (C) Quantification of molecular expression in P-EN and P-FN neurons. P-EN, n = 13; P-FN, n = 6 brains. In all panels, LexA+ neurons in green, Runt in magenta, and Cut in cyan. Yellow outline, neurons of interest. Scale bars: 10 μm.

### Loss of Svp decreases the number of late born P-FN adult neurons

Svp is required for the early-to-late switch in T2NB temporal gene expression (Ren et al., 2017; Syed et al., 2017). Here, we test whether this Svp-dependent switch in the T2NB gene expression extends to the specification of post-mitotic neurons. In this section we asked whether Svp knockout reduces late born P-FN neurons. We found Svp knockout leads to a highly penetrant loss of late born adult P-FN neurons (Fig. 4A-D). We note that there are some P-FN neurons remaining; because the cell bodies are closely clustered, they likely reflect that the Svp knockout failed to remove *svp* within individual T2NB lineages (see Discussion). Further evidence that remaining P-FN neurons derived from Svp escaper NBs include their normal morphology (Fig. 4E-I; S. Movies 1-2) and normal birth date (Fig. 4J). We conclude that Svp is required for the T2NBs to produce the late born P-FN neuron identity, likely due to an extension of early born neuron identities (Fig. 4K).

**Figure 4.**
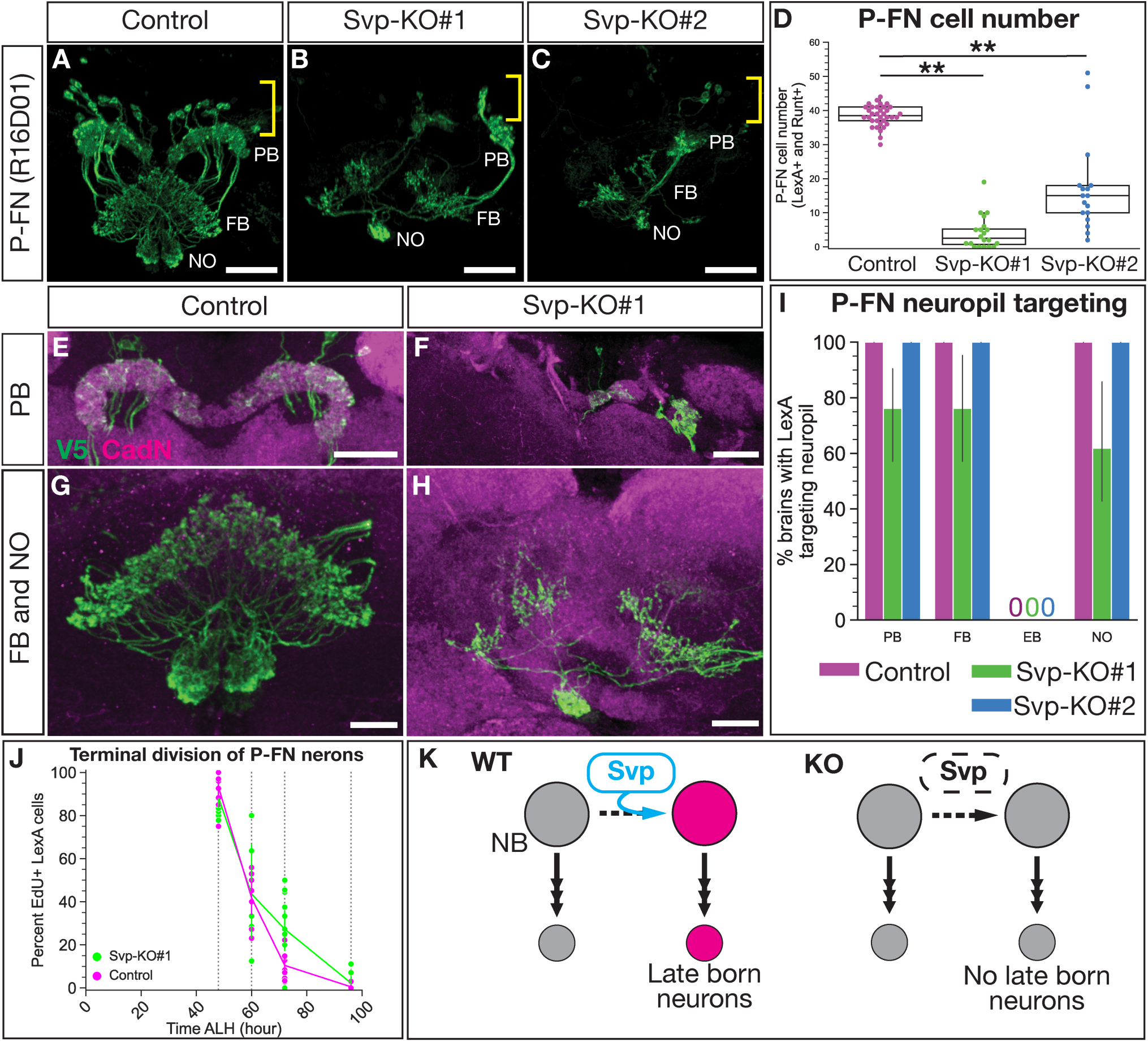
Svp is required for the late born P-FN neuron identity. (A-C) P-FN neurons labeled by LexA driver expressing membrane bound V5 show loss of Svp leads to the loss of P-FN neurons. Yellow brackets show cell body region and white labels of neuropils: Protocerebral bridge (PB), fan-shaped body (FB), and noduli (NO). (D) Quantification of P-FN cell number identified with co-expression of LexA and Runt. Each dot represents one adult brain with box and whisker plot showing distribution. Control, n = 34; Svp-KO#1, n = 20; Svp-KO#2, n = 17. *P*-values determined by One-way ANOVA, *P* < 0.001, with Tukey post-hoc test: Control versus Svp-KO#1 *P* < 0.001, Control versus Svp-KO#2 *P* < 0.001. (E-F) P-FN neuron PB morphology shows lack of targeting with loss of Svp. (G-H) P-FN neuron FB and NO morphology shows lack of targeting with loss of Svp. (I) Quantification of P-FN neuropil targeting scored based on LexA targeting to neuropils identified with nc82 or CadN shown as a bar plot with 95% confidence interval. Control, n = 35; Svp-KO#1, n = 21; Svp-KO#2, n = 13; 95% confidence interval. (J) P-FN EdU dropout shows P-FN neurons born for Svp-KO escaper neuroblasts are born in a normal NB birth window as shown by percent P-FN neurons labeled by EdU. Each dot represents one adult brain. For all timepoints, Control, n = 11-12; Svp-KO#1 = 5-12 brains. (K) Summary of Svp required for P-FN identity. In all images, LexA+ neurons driving V5 in green and CadN in magenta. Scale bars: (A-C) 20 μm, (E) 30 μm, (F) 20 μm, (G-H) 10 μm.

### Loss of Svp extends the production of early born P-EN adult neurons

We next tested whether Svp knockout in T2NBs extends early born P-EN neuron identity, using both molecular and morphological assays. We found that loss of Svp leads to the expansion of adult P-EN neurons, with projections into the protocerebral bridge, ellipsoid body, and noduli (Fig. 5A-C). Furthermore, these ectopic P-EN neurons expressed the appropriate P-EN molecular markers: P-EN LexA-driven V5, Runt, and Cut (Fig. 5D-F’’’,G). We conclude that Svp is required to restrict the production of P-EN neurons.

**Figure 5.**
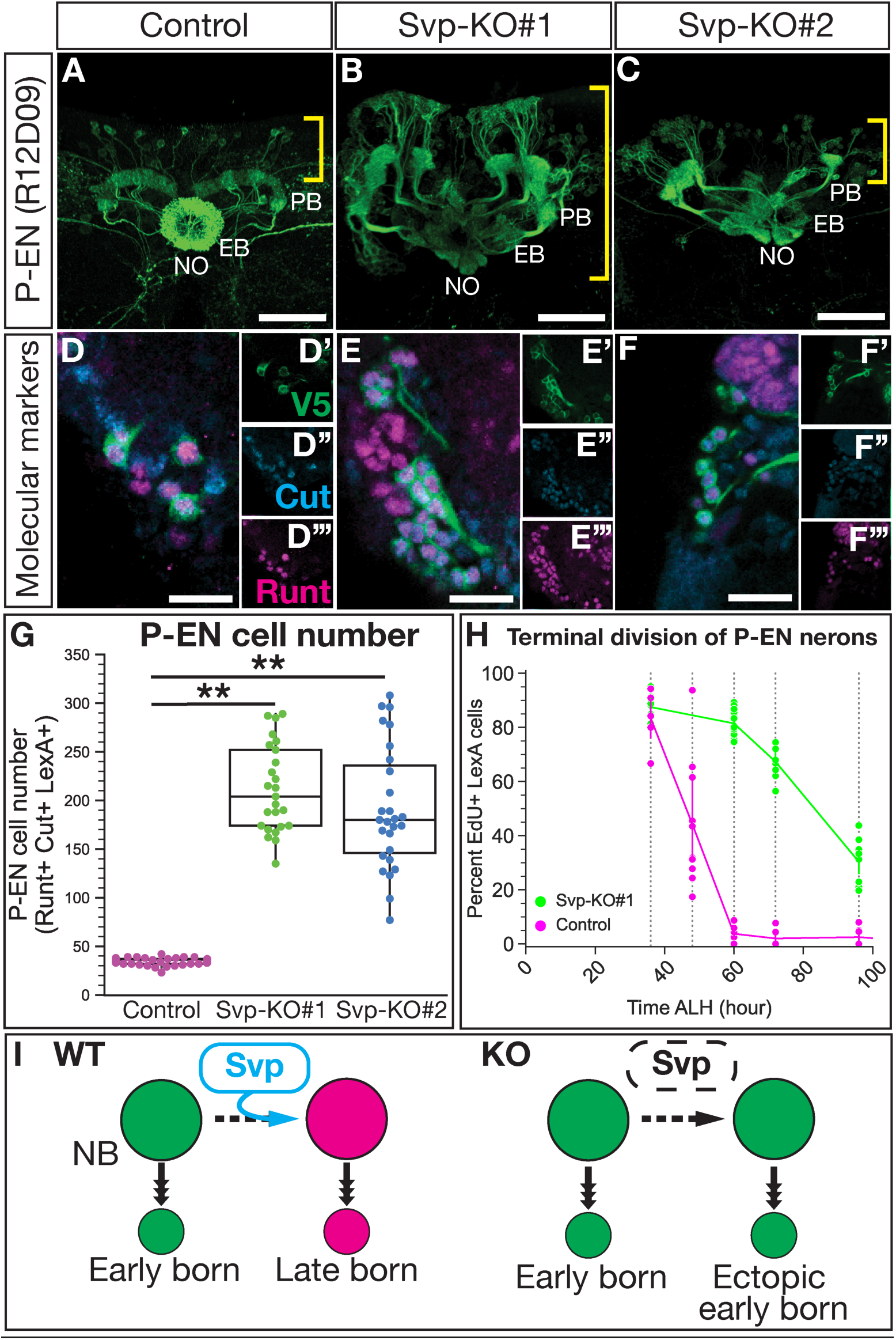
Svp restricts the early born P-EN neuron molecular identity and birth window. (A-C) P-FN neurons labeled by LexA drivers show loss of Svp produces ectopic P-EN neurons. Yellow brackets show cell body region and white labels of neuropils: Protocerebral bridge (PB), ellipsoid-body (EB), and noduli (NO). (D-F’’’) Ectopic neurons maintain expression of P-EN LexA, Cut and Runt. (G) Quantification of P-EN cell numbers from A-F. Each dot represents one adult brain with box and whisker plot showing distribution. Control, n = 35; Svp-KO#1, n = 25; Svp-KO#2, n = 27 brains. *P*-values determined by One-way ANOVA, *P* < 0.001, with Tukey post-hoc test: Control versus Svp-KO#1 *P* < 0.001, Control versus Svp-KO#2 *P* < 0.001. (H) EdU dropout of P-EN neurons show an extended birth window with loss of Svp shown by percent P-EN neurons labeled by EdU. Each dot represents one adult brain. Control, n = 6-10; Svp-KO#1, n = 8-13. (I) Summary of Svp restricting P-EN neuron molecular identity and birth window. In all images, LexA+ neurons driving V5 in green, Runt in magenta, and Cut in cyan. Scale bars: (A-C) 40 μm, (D-F’’’) 10 μm.

The ectopic P-EN neurons could arise from T2NBs generating more P-EN neurons at a normal early time, or they could arise from T2NBs generating P-EN neurons at abnormally late times in their lineage (replacing late born P-FN neurons). To distinguish these models, we performed EdU birth dating of P-EN neurons following Svp knockout. Whereas wild-type P-EN neurons are all born prior to 60h ALH (Fig. 1B), we found that Svp knockout resulted in P-EN neurons still being born at 100h ALH (Fig. 5H). Our data supports a model in which loss of Svp extends the production of early born P-EN neurons at the expense of late born P-FN neurons (Fig. 5I).

We next wanted to know if the ectopic P-EN neurons showed normal neuropil targeting. We reconstructed CX neuropils from anti-nc82 (Bruchpilot; a presynaptic marker) (Wagh et al., 2006) stains and assayed P-EN neuropil targeting (Fig. 6; S. Movies 3-4). Note that we first quantified all CX neuropil volumes, which includes more neuron subtypes than just the P-EN and P-FN neurons. We found that following loss of Svp in the T2NB lineages, all CX neuropils increased in size (Fig. S4A). This is due in part to ectopic P-EN neurons, which targeted all assayed CX neuropils but with increased targeting volume (Fig. 6A-H’; Fig. S4B-C). The role of neurons in addition to P-EN in generating this neuropil enlargement phenotype is unknown. Next, we assayed neuropil targeting of ectopic P-EN neurons in our T2NB Svp knockouts. We found that ectopic P-EN neurons target all the expected CX neuropils (e.g., protocerebral bridge, ellipsoid body, noduli) but with increased volume, likely reflecting the increase in neurons (Fig. 6A-F’; Fig. S4B). Surprisingly, we observed ectopic P-EN neurons abnormally target the fan-shaped body (Fig. 5G-H’,I, Fig. S4B). We conclude that loss of Svp leads to the generation of ectopic P-EN neurons which show targeting to normal neuropil regions with increased volume and off-targeting to the fan-shaped body.

**Figure 6.**
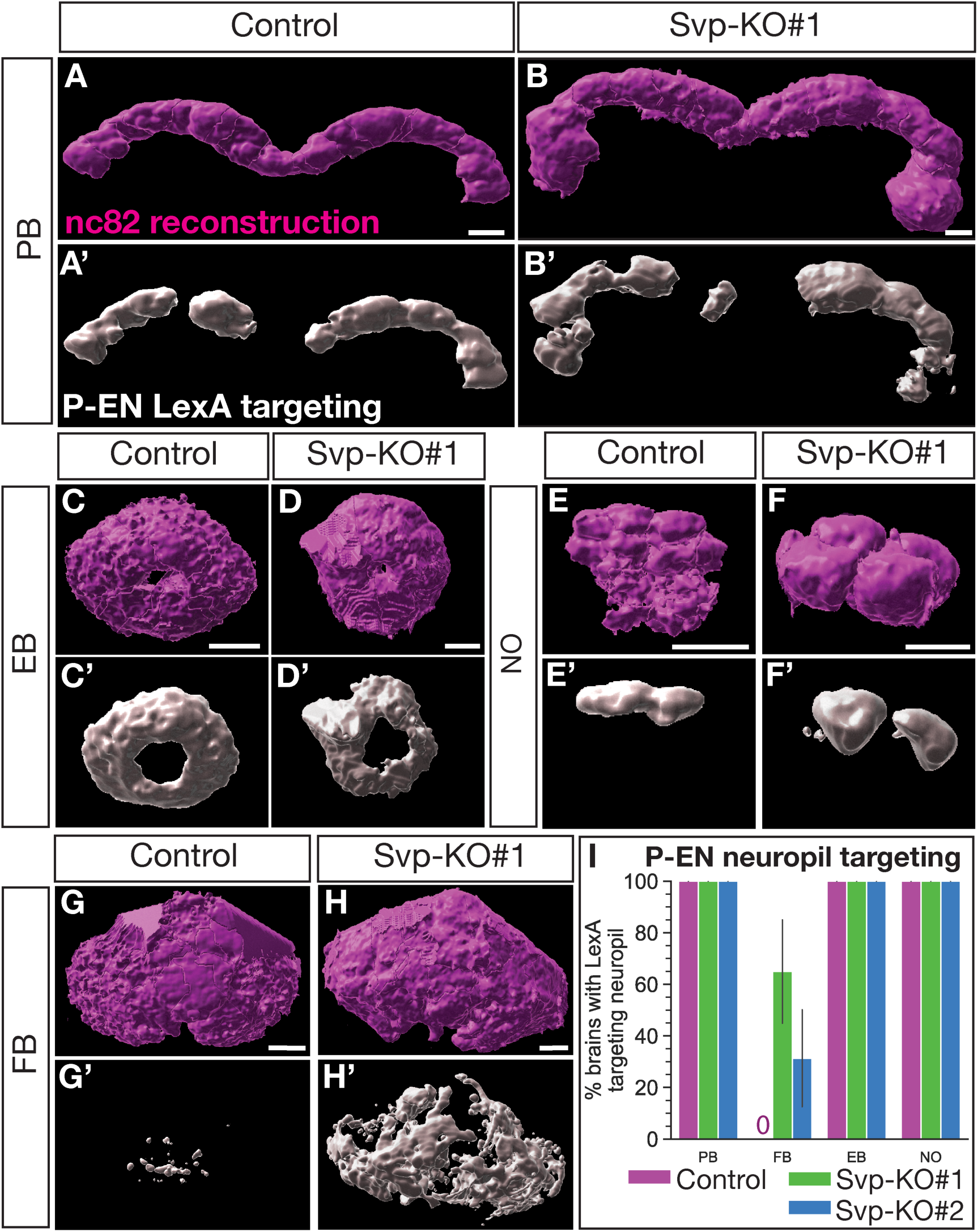
Svp regulates the early born P-EN adult neuron morphology. (A-H’) Reconstruction of CX neuropils and P-EN neuron targeting for the protocerebral bridge (PB; A-B’), ellipsoid body (EB; C-D’), noduli (NO; E-F’), and fan-shaped body (FB; G-H’). (I) Quantification of P-EN neuropil targeting scored based on LexA targeting to neuropils identified with nc82 or CadN shown as a bar plot with 95% confidence interval. Control, n = 8; Svp-KO#1, n = 20; Svp-KO#2, n = 16 brains. In all panels, nc82 reconstruction in magenta and P-EN LexA targeting in white. Scale bars: 10 μm.

### Loss of Svp extends T2NB lineages into the adult

We note above that Svp knockout extended the production of P-EN neurons beyond the later part of larval life at 100h ALH (Fig. 5H), raising the question: Is Svp required for terminating neurogenesis in T2NBs? In wild-type animals, most NBs undergo decommissioning -- characterized by loss of molecular markers, cell cycle arrest, death and/or differentiation -- in late larval or early pupal stages (Ito and Hotta, 1992; Maurange et al., 2008; Siegrist et al., 2010; Yang et al., 2017). The time of T2NB decommissioning has not been previously reported. We found that T2NBs normally complete decommissioning by 24h after pupal formation (APF) (Fig. 7A-A’’’,E). In contrast, Svp knockout in T2NB lineages results in delayed decommissioning with persistence of proliferative T2NBs, marked by pH3, into 7-day old adults (Fig. 7B-D’’’,E-F). Remarkably, the persistent T2NBs following Svp knockout strongly express the early marker Imp and lack the late marker Syncrip (Fig. 7B-D’’), consistent with previous work showing the Imp to Syncrip gradient regulates NB decommissioning (Yang et al., 2017). Thus, we conclude that Svp is required to initiate T2NB decommissioning (Fig. 7G). How these late functions of Svp, such as T2NB decommissioning in pupal stages, are triggered by transient expression of Svp many days earlier in first instar larvae remains an interesting open question.

**Figure 7.**
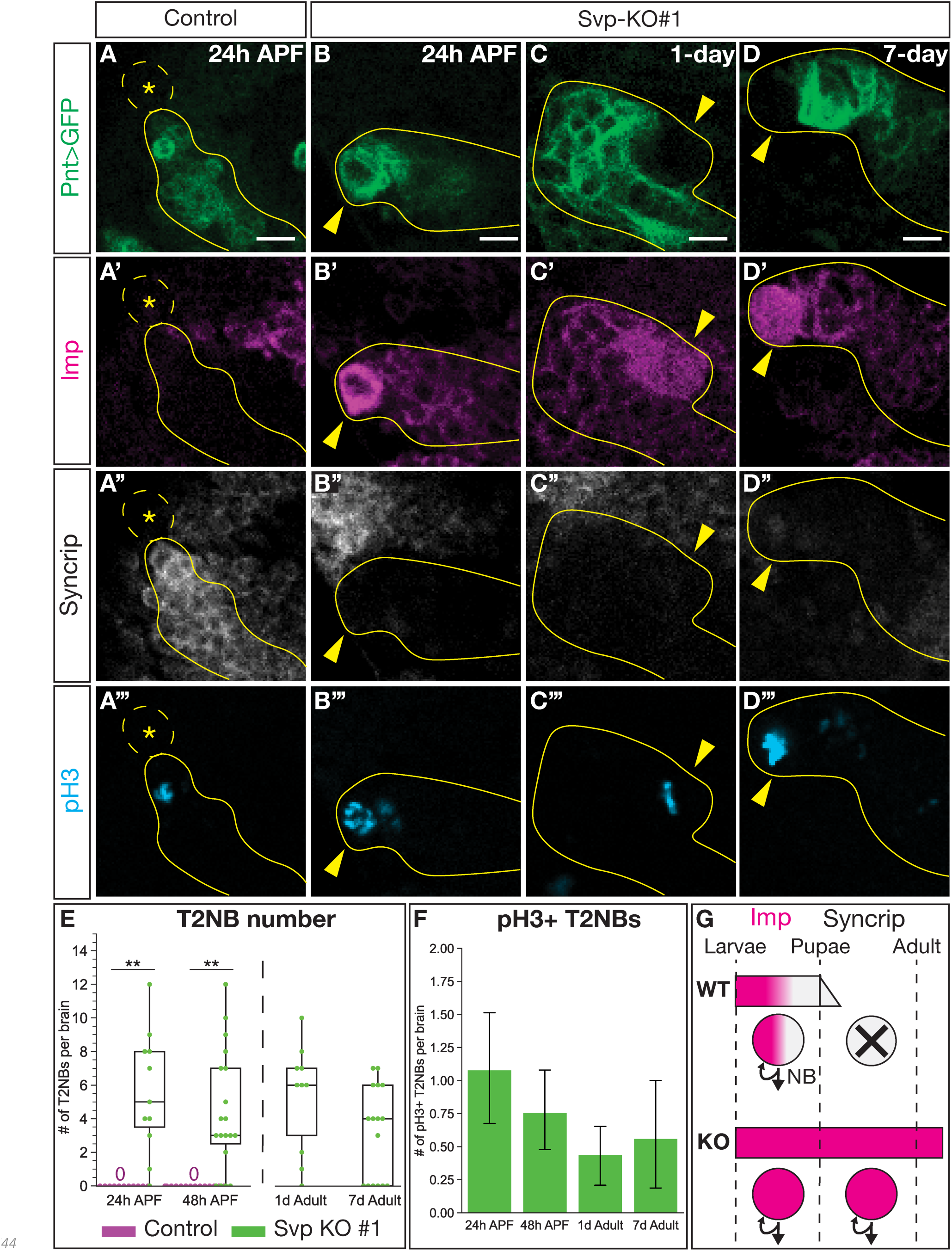
Svp is required for timely onset of T2NB decommissioning. (A-A’’’) T2NBs have decommissioned by 24h after pupal formation (APF). (B-D’’’) Loss of Svp lead to T2NBs remaining with expression of the early factor Imp and no expression of the late factor Syncrip while being mitotically active with expression of pH3 at 24h APF (B-B’’’), 1-day adult (C-C’’’) and 7-day adult (D-D’’’). (A-D) In all images, Type 2 lineage identified with Pnt-Gal4 in green, Imp in magenta, Syncrip in white, and pH3 in cyan. Dashed outline and asterisk, lack of T2NB. Solid yellow outline, Type 2 lineage. Yellow arrowhead, T2NB. (E) Quantification of T2NB number per brain. Each dot represents one brain with box and whisker plot showing distribution. Control 24h APF, n = 16; Svp-KO#1 24h APF, n = 11; Control 48h APF, n = 18; Svp-KO#1 48h APF, n = 19; Svp-KO#1 1d adult, n = 10; Svp-KO#1 7d adult = 16. *P*-values determined by an independent t-test: Control versus Svp-KO#1 at 24h APF *P* < 0.001, at 48h APF *P* < 0.001. (F) Quantification for number of pH3+ ectopic T2NBs remaining in the pupae and adult stages shown as a bar plot with 95% confidence interval. Svp-KO#1 24h APF, n = 37; Svp-KO#1 48h APF n = 25; Svp-KO#1 1d adult, n = 26; Svp-KO#1 7d adult, n = 16. (G) Summary of Svp required for transition of Imp to Syncrip expression in T2NBs for onset of neuroblast decommission. Scale bars: 5 μm.

## Discussion

### Columnar neurons are born at different times in the T2NB lineage

Our EdU birth dating revealed that P-EN neurons are born from early T2NBs, and P-FN neurons are born from late T2NBs. Previous work showed these neuron identities are both born during the same NB window, conflicting with our results (Sullivan et al., 2017). Our approach tracks the terminal division of cells through EdU labeling, which accounts for all neurons within the population, while previous birth dating relied on a genetic approach that only accounted for 55-65% of the neurons (Sullivan et al., 2017). We propose that EdU birth dating is more comprehensive for assigning neurons to NB birth windows as all neurons are reliably labeled with EdU, providing reproducible tracing of terminal divisions.

It remains an open question how T2NBs are patterned to generate birth-order dependent neuronal identities. One hypothesis is that the lineage uses a temporal transcription factor cascade, similar to embryonic Type 1 NBs (Isshiki et al., 2001; Novotny et al., 2002; Pearson and Doe 2003; Tran and Doe 2008; Moris-Sanz et al., 2014). Alternatively, or in combination, the lineage could utilize a temporal gradient of protein expression, like mushroom body NBs (Liu et al., 2015). Several temporally expressed factors are known to cross-regulate in T2NBs (Ren et al., 2017; Syed et al., 2017). Our work with loss of Svp (i.e., changing expression of known temporal factors in T2NBs) shows that temporal patterning at the NB level is required to specify neuron subtypes. This is supported by recent work reporting knockdown of Imp levels in T2NB lineages results in altered CX neurons; however, it is unclear if this is due to the role of Imp at the NB or INP level (Hamid et al., 2023). Additionally, loss of T2NB temporal factors EcR and E93 are required for specifying the CX neuron subtype dFB neurons (Wani et al., 2023). Our data do not distinguish between a model for a temporal transcription factor cascade and/or temporal gradients. Thus, it will be important to test individual temporal factors and expression levels in specifying birth-order dependent fates.

### Svp expression in larval T2NB lineages

We report a comprehensive characterization of Svp in all eight T2NB lineages at stages that bracket Svp expression. Previous work has only observed Svp expression at either broad temporal windows or limited to a few lineages (Bayraktar and Doe, 2013; Ren et al., 2017; Syed et al., 2017). We show that Svp has a tight expression window between 18-24h ALH in all T2NB lineages with Svp protein restricted to the NB (Fig. 2). This finding is further supported by the *Svp* mRNA showing a similar expression pattern in the T2NB lineages and restricted to the NB (Fig. S1). Interestingly, *svp* mRNA was expressed before onset of protein expression and remained longer than protein expression (compare Figure 2 to Sup Fig. 1), which suggests post-transcriptional regulation of *Svp* in the NB. We found evidence that indicates the posterior lineages (DM4-6, DL1-2) express Svp prior to the anterior lineages (DM1-3) (Fig. 2I). We speculate that the distinct onset of Svp expression between lineages may be determined by differential expression of spatial factors. Determining the upstream mechanism that initiates Svp expression remains an open question. A further challenge will be resolving the orphan receptor status of Svp as it remains unknown if Svp requires a ligand for activation.

### Svp is required for late born fates in the T2NB lineages

We found that Svp knockout resulted in a loss of the late born P-FN neuron identity. The remaining P-FN neurons resemble wild-type neurons in cell body location, numbers, morphology, and birth window. We report that the Svp knockout lineages have a success rate of ∼75%, with approximately 1 out of 4 T2NBs completely escaping *svp* knockout (Fig. S2). Wild-type P-FN neurons number ∼40 cells, but in Svp knockouts we see ∼5-15 cells remaining, which is the expected proportion based on the efficiency of our knockout lines (Fig. 4D). Moreover, the remaining P-FN neurons display lineage-appropriate morphology (Fig. 4E-H) and a wild-type birth window from the T2NBs (Fig. 4J). We conclude that the remaining P-FN neurons are derived from Svp knockout escaping NBs that maintain normal Svp expression and hence develop normally. Thus, our Svp knockout can be described as an “all or none” knockout of Svp in ∼75% of the T2NBs.

### Svp specification of other CX neuron subtypes

Svp has previously been reported as either a switching factor in embryonic and larval NBs (Kanai et al., 2005; Mettler et al., 2006; Maurange et al., 2008; Benito-Sipos et al., 2011; Kowhi et al., 2011) or as a post-mitotic fate determinate in photoreceptor neuron subtypes (Mlodzik et al., 1990; Hiromi et al., 1993). Our work shows that Svp acts as a switching factor in T2NBs for specifying neuron identities, as Svp was not expressed in post-mitotic P-EN or P-FN neurons (Fig. S3). However, we cannot rule out that Svp is required as a post-mitotic fate determinant for other CX subtypes.

We show that Svp is required to restrict the early born P-EN identity as loss of Svp resulted in T2NBs extending production of P-EN neurons into an abnormally late temporal window (Fig. 5H). These ectopic P-EN neurons maintain wild-type molecular markers (e.g., LexA, Runt, Cut) (Fig. 5G). We note that these ectopic P-EN neurons have ectopic targeting to the fan-shaped body, a neuropil not targeted by wild-type P-EN neurons (Fig. 6G-H’,I; Fig. S4B-C). We speculate that this abnormal targeting may be compensation for the loss of P-FN neurons that normally target the region, suggesting that columnar neuron targeting may be promiscuous when subtypes are absent. We note that P-EN neurons are just one fate born from the early temporal window; other neurons born at a similar time are not visible with the P-EN/P-FN LexA markers used here. We hypothesize that these early born populations will also be expanded with the loss of Svp. It will be vital for future work to identify additional CX neuron markers and their birth date from the T2NBs to test for functional temporal patterning factors.

We find that the loss of Svp resulted in significant changes to CX neuropil volumes. While targeting from ectopic P-EN neurons accounts for a portion of this increase, it does not fully account for the global CX neuropil enlargement we report (Fig. S4B-C). We speculate that either expanded populations of other T2NB-derived neuron subtypes and/or synaptic partners account for the ectopic targeting to these neuropils. For example, P-EN neurons form synapses with the T2NB-derived E-PG neurons (Green et al., 2017; 2019), thus, loss of Svp could lead to either E-PG compensation with increased targeting to ectopic P-EN neurons and/or an expanded E-PG neuron population. We are unable to account for the global CX changes that occurred with the loss of Svp due to the limitation in our assays of just the P-EN and P-FN neurons.

### Svp regulation of Type 2 neuroblast temporal progression

Svp is expressed in early larval T2NBs prior to the Imp to Syncrip transition (Ren et al., 2017); thus, we were surprised by our findings that Svp is required for terminating neurogenesis in pupae. Previous work has shown the transition in central brain NBs from Imp to Syncrip expression initiates NB decommissioning (Yang et al., 2017). Interestingly, mushroom body NBs remain proliferating into pupal stages, when other NBs have decommissioned, due to sustained Imp expression (Yang et al., 2017). Consistent with our work on T2NBs, previous work has also demonstrated that loss of Svp in ventral nerve cord and central brain NB lineages leads to continued NB proliferation into the adult (Maurange et al., 2008; Narbonne-Reveau et al., 2016). We report that loss of Svp in early larval T2NBs had profound impacts on the lifespan of the NBs, where they survive into 7-day adults manifesting (i) maintained Imp expression; (ii) mitotically active; (iii) maintain low levels of Syncrip (Fig. 7). Thus, we propose that Svp initiating the switch of Imp-to-Syncrip in T2NBs is a NB autonomous mechanism required for terminating neurogenesis within T2NB lineages. This is consistent with previous work showing Svp is required for the progression of Imp to Syncrip expression in T2NBs (Ren et al., 2017). We speculate that Svp may regulate chromatin landscape and/or initiate a temporal cascade to account for the long-term effects Svp has on T2NB temporal progression.

Recently, Notch signaling has been shown to be required for central brain Type 1 NB decommissioning by disrupting temporal patterning progression; loss of Notch signaling produced prolonged expression of the early factor Imp and reduced expression of the late factor E93 (Sood et al., 2023). Additionally, Notch signaling appeared to be required in terminating expression of the early factors Castor and Svp (Sood et al., 2023). This indicates that Notch signaling acts upstream of NB temporal factors, and thus is likely to act upstream of Svp as well. This is difficult to test as loss of Notch signaling in T2NBs results in loss of neuroblast identity (San-Juan and Baonza, 2011; Zhu et al., 2012; Li et al., 2016; Li et al., 2017).

### Conserved role of Svp in vertebrate temporal patterning

Our work shows that Svp acts as a switching factor in T2NBs to switch from producing early born to late born neuronal identities. The mammalian orthologs of *svp*, *Coup-tfI/II* have been characterized for a similar role as a switching factor in murine neural stem cells (Naka et al., 2008; Lodato et., al 2011). *Coup-tfI/II* are required for cultured neural stem cells to switch from producing early born neuron cell types to producing late born glia, since a loss of this molecular switch resulted in sustained neurogenesis (Naka et al., 2008). Additionally, *Coup-tfI/II* are required for switching cortical neural stem cells from producing early born interneuron fates to late born interneuron fates (Lodato et., al 2011). These findings, along with ours and others in *Drosophila*, suggest that Svp has a conserved role as a neural stem cell switching factor from fly to mammals (Mettler et al., 2006; Benito-Sipos et al., 2011; Ren et al., 2017).

## Materials and Methods

### Animal Preparation

*Drosophila melanogaster* was used in all experiments. All flies were kept and maintained at 25°C unless stated otherwise. Stocks used can be found in Table S1 and experimental genetic crosses in Table S2.

#### EdU experiments

EDU (5-ethynyl-2’-deoxyuridine; Millipore-sigma #900584-50MG), a thymidine analog, was used to label proliferating cells starting at various sequential larvae ages. Larvae were fed food containing 20ug/ml EdU nonstop from the initial age feeding started until the larvae pupated. Larvae fed on EdU were raised at temperatures between 18°C to 21°C until adults hatched and were dissected.

#### Larval experiments

Embryos were collected on 3% agar apple juice caps with yeast paste for 4 hours and aged for 21 hours. After aging, embryos were transferred to a fresh cap and aged 4 hours for hatching. Hatched larvae were collected and dissected at the corresponding time after larval hatching (ALH).

#### Adult experiments

Males and virgin females were introduced in standard yeast medium vials and flipped every two days. 2-5 day old adult flies were dissected for all experiments unless stated otherwise. All animals dissected were a mixture of male and female unless otherwise specified.

### Hybridization Chain Reaction (HCR) RNA fluorescent in situ hybridization

Larval brains were dissected in Schneider’s insect medium, fixed in 4% PFA (paraformaldehyde; Electron Microscopy Sciences 15710) in PBS (Phosphate buffered saline, Sigma-Aldrich P4417) for 7-15 min at room temperature, and washed in PBST (PBS with 0.3% Triton, Sigma-Aldrich T8787). The fixed brains were stored in 70% Ethanol in water at 4 °C until used. We followed the protocol from Duckhorn et al. (2022). A 20-probe set targeting svp transcripts was synthesized by Molecular Instruments, Inc. and probes were added to a final concentration of 4 nM for hybridization. Amplifier B3-546 was also synthesized by Molecular Instruments, Inc and 6 pmol of each hairpin (h1 and h2), was added for amplification.

### Immunohistochemistry

Antibodies used with supporting notes can found in Table S3.

#### Larval brain sample preparation

Larval brains were dissected in PBS and mounted on poly-D-lysine coated coverslips (Neuvitro Corporation GG-12-PDL; primed in 100% ethanol). Samples fixed for 23 minutes in 4% PFA in PBST. Samples were washed in PBST and blocked with 2% normal donkey serum (Jackson ImmunoResearch Laboratories, Inc.) in PBST. Samples incubated in a primary antibody mix diluted in PBST for overnight or 1-2 days at 4°C. Primary antibodies were removed, and samples thoroughly washed with PBST. Samples were incubated in secondary antibodies overnight at 4°C. Secondary antibodies were removed, and samples washed in PBST. Samples were dehydrated with an ethanol series of 30%, 50%, 75%, and 100% ethanol then incubated in xylene (Fisher Chemical X5-1) for 2×10 minutes. Samples were mounted onto slides with DPX (Sigma-Aldrich 06552) and cured for 3-4 days then stored at 4°C until imaged.

#### Adult brain sample preparation

Adult brains were prepared similar to larval brains with the exception of 41 minutes for fixation in 4% PFA and 2×12 minute xylene incubations.

#### EdU adult brain sample preparation

Adult brains from EdU fed larvae were dissected in HL3.1 then fixed in 4% PFA for 30 minutes and incubated in block at 4°C overnight. Samples were incubated in primary and secondary mixes prior to Click-it-Reaction to label EdU. The Click-it-Reaction mix was comprised of PBS, Copper II sulfate (ThermoFisher 033308.22), 555-Azide (Thermofisher A20012) in DSMO and ascorbic acid (Sigma-Aldrich A4544-25G) for a 2-hour incubation. Samples were dehydrated and washed in xylene before DPX mounted as described above.

### Confocal Microscopy

Fixed preparations were imaged with a Zeiss LSM 900 or 800 laser scanning confocal (Carl Zeiss AG, Oberkochen, Germany) equipped with an Axio Imager.Z2 microscope. A 10x/0.3 EC Plan-Neofluar M27 or 40x/1.40 NA Oil Plan-Apochromat DIC M27 objective lens were used. Software program used was Zen 3.6 (blue edition) (Carl Zeiss AG, Oberkochen, Germany).

### Image processing and analysis

#### Cell counting and neuropil target scoring

Confocal image stacks were loaded into FIJI (ImageJ 1.50d, https://imagej.net/Fiji). Cells were counted using the Cell Counter plugin. Neuropil targeting was determined by co-localization of LexA expression with neuropil marker that was not a filament bundle passing through the neuropil.

#### Imaris neuropil reconstructions

Confocal image stacks were loaded into Imaris 10.0.0 (Bitplane AG, Zurich, Switzerland). Imaris Surface objects were created for each neuropil using nc82 staining and LexA expression followed by new objects designating overlap between neuropil and neurites. Briefly, the Surface tool was selected, and a region of interest (ROI) was drawn to encompass a whole CX neuropil or LexA expression. The source channel was selected (nc82 in RRX for neuropils or LexA in 647 for neurites) and absolute threshold intensity was manually set slice by slice to outline fluorescent signal and morphologically split to separate regions. All Starting Points and Seed Points were kept ensuring full coverage of signal. Surfaces were rendered and surfaces outside neuropil structures removed. To find the LexA targeting for each neuropil, Surface-Surface Overlap File XTension (Matthew Gastinger, https://imaris.oxinst.com/oP-EN/view/surface-surface-overlap) was used to find the volume (μm^3^) of overlap. A Smoothing Factor of 0.2 μm was kept for all surfaces.

#### Figure preparation

Images in figures were prepared either in Imaris 10.0.0 or FIJI. Scale bars are given for a single slice in all single slice images and from all stacks within maximum intensity projections images. Pixel brightness was adjusted in images for clearer visualization; all adjustments were made uniformly over the entire image, and uniformly across wild-type samples and corresponding control and experimental samples. Adobe Illustrator 2023 was used for formatting.

### Statistical analyses

Statistics were computed using Python tests (see supplemental script for specific packages). All statistical tests used are listed in the figure legends. *P*-values are reported in the figure legends. Plots display n.s. = not significant, * is *P* < 0.05, and ** is *P* < 0.01. Plots were generated using Seaborn and Matplotlib packages in Python.

## Additional files

<u>Supplementary figures</u>

<u>Supplementary tables</u>

## Acknowledgements

We thank fellow lab members Kristen Lee, Peter Newstein, Megan Radler, and Chundi Xu for constructive comments on the manuscript. We also thank Tzumin Lee (University of Michigan), and Mubarak Syed (University of New Mexico) for comments on the manuscript. Antibodies obtained from the Developmental Studies Hybridoma Bank, created by the NICHD of the NIH and maintained at the University of Iowa, Department of Biology, Iowa City, IA were used in this study. Stocks obtained from the Bloomington Drosophila Stock Center (NIH P40OD018537) and Vienna Drosophila Resource Center were used in this study.

## Author contributions

Conceptualization: NRD, CQD. Design: NRD, LM (EdU drop out), KH (HCR RNA in situ), CQD. Investigation: NRD, LM (EdU drop out), KH (HCR RNA in situ). Analysis: NRD. Writing – original draft: NRD, CQD. Writing – review and editing: NRD, LM, KH, CQD.

## Data availability

All relevant data can be found within the article and its supplementary information.

